# Population variation in lipid depletion during parental care in male threespine sticklebacks (*Gasterosteus aculeatus*)

**DOI:** 10.1101/2022.12.16.520737

**Authors:** Robert Scott

## Abstract

I examined changes in stored lipid over the parental care interval in several populations of threespine stickleback (*Gasterosteus aculeatus*). Males from the limnetic populations and the marine population decreased in stored lipids over the progression of the nesting interval. Of the two benthic populations, one (Hotel Lake) did not decrease stored lipids over the parental care interval, as predicted, whereas the other (Crystal Lake) showed declining stored lipids over the parental care interval. Though food resources are likely high in the nesting area of Crystal Lake, the high density of conspecific cannibal groups in the littoral zone likely contributes to reduced foraging and elevated guarding/diversionary display behaviour and dependency on stored lipids to fuel those behaviours in the Crystal Lake (benthic) population. This study demonstrates the value of considering different ecological conditions on behavioural and energetic constraints among populations of threespine stickleback.

## Introduction

Males of several teleost fish undertake extended, sole parental care (Wooton 1989). Activities associated with parental care are energetically intensive and maintaining such intensive activities is often dependent on stored energy reserves since foraging opportunities can be limited. For example, in smallmouth bass males undertake parental care that can last for several weeks, and although most males forage to a limited extent, all deplete lipid reserves during the parental care interval (Gravel and Cooke 2013; Mackereth et al. 1993). Feeding during parental care activities offsets physiological stress associated with parental care and impacts reproductive success positively (Zolderdo et al. 2016) and may reduce occurrence of filial cannibalism when energy reserves become depleted (Manica 2002).

Male threespine stickleback (*Gasterosteus aculeatus*) exhibit extended parental care lasting up to 2 weeks (Whoriskey and FitzGerald 1994). During the reproductive interval, males move into shallow (0.5 – 3 m) areas along the shore, establish an exclusive territory (0.5 – 2 m^2^), construct a tube-shaped nest using a variety of materials held together with kidney secretions, and then court females. Once a male has a clutch of eggs (from one or more females) he will begin to fan the developing embryos and chase away territorial intruders, some of which are likely predators (eg. Foster 1993; Scott and Foster 2000). Males in some populations continue to guard their free-swimming offspring following their emergence from the nest (Whoriskey and FitzGerald 1994).

Foraging opportunities for nesting male sticklebacks may be high since they nest in shallow, littoral zones in lakes so their dependence on stored energy reserves during parental care is not an issue (Wooton 1984). However, males have been observed to lose weigh and decrease stored lipids over the parental care interval (Chellappa et al. 1989; Dufresne et al. 1990) suggesting that they are dependent on stored energy reserves at least to some extent.

Threespine stickleback populations vary in prey specialization; some population are characterized as benthic foragers (adapted to feed on benthic prey) whereas others are characterized as limnetic foragers (specialized for foraging on planktonic prey) (McPhail 1984; Schluter 2000). In benthic populations, nesting males tend to interact with non-neighbours but limnetic males interact primarily with neighbours (Foster et al. 2008). This observation suggests that there is abundant food in the nesting area in lakes with benthic types (ie., lots of individuals are in the nesting area to forage on abundant food), but that food is not abundant in the nesting area in lakes with the limnetic type (ie few non-nesting individuals are there foraging). Schluter (1995) showed that limnetic and benthic feeding specialists grew slower when provided with the alternative food sources; benthic specialists grew slower when forced to forage on limnetic prey and limnetic specialist grew slower when forced to forage on benthic prey compared to each foraging on the prey to which they were specialized. Therefore, limnetic types are probably limited in their foraging opportunities during nesting compared to benthic types in their respective lakes, and more dependent upon stored energy reserves during parental care than their benthic counterparts. I collected nesting male threespine stickleback from each of two limnetic and two benthic freshwater populations to compare lipid changes over the reproductive interval. I predicted that parental males from benthic populations would not deplete lipid stores over the reproductive interval whereas males from limnetic populations would show a decrease in lipid stores. I also collected nesting males from a marine location since marien populations represent the ancestral state for both benthic and limnetic types (Bell and foster 1993).

## Methods

I collected nesting male stickleback from five populations in British Columbia, Canada, two representing limnetic populations (Cowichan Lake and Garden Bay Lake), two representing Limnetic populations (Hotel Lake and Crystal Lake) and one representing a marine population (Swy-a-Lana Lagoon; figure 1) (Foster 1994; Foster 2013; Foster et al. 2019). Individual nesting males were located (using mask and snorkel) and observed until their stage of reproduction (courting, caring for embryos in the nest - early parental care, or caring for free-swimming offspring - late parental care phase) could be determined. Nesting males were subsequently captured using a dipnet and transported to shore where they were euthanized and preserved individually on dry ice. Samples were transferred to a freezer (−80°C) and stored prior to processing for lipid analysis.

**Figure 1.**
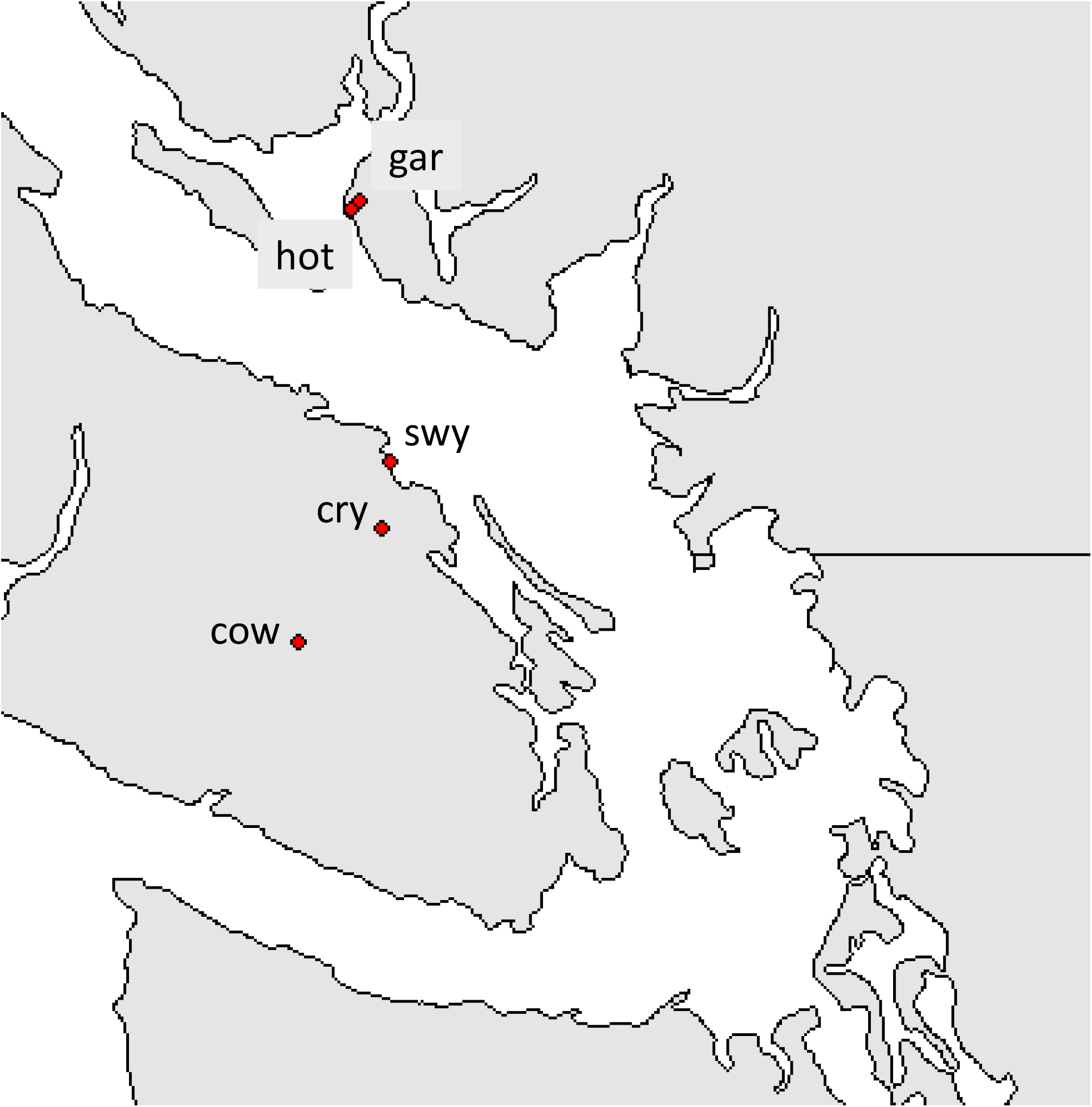
Map of southern British Columbia showing the location of each sampled population (cow – Cowichan Lake; cry – Crystal Lake; swy – Swy-a-Lana Lagoon; gar – Garden Bay Lake; hot – Hotel Lake).

Individual samples were thawed and their length (to the nearest 0.01mm) and whole-body mass (nearest 0.0001 g) were measured. The left musculature and dermis of each individual was removed, ground to a powder using a frozen mortar and pestle and transferred to a pre-weighed envelope. Samples were subsequently dried at 75°C for 72 h and lipids were extracted using a Soxhlet apparatus and petroleum ether (boiling point 35-60°C) for 20 h. Samples were air dried for 8 h and subsequently for 12 h at 75°C following extraction. Each sample was weighed before and after each step of the process. Sample lipid was determined by subtracting the post-Soxhelt weight rom the pre-Soxhlet extraction weight and tissue concentration was calculated as g lipid * g sample dryweight^-1^.

Statistical analyses were performed using the RStudio environment (version 2021.09.2+382; R Studio Team 2020) for R (version 4.1.3; R Core Team 2022). We adjusted for the effect of body weight on lipid concentration following the method outlined by Lleonart et al. (2000). Size adjusted lipid concentration for each individual across all linear traits were calculated using the following:

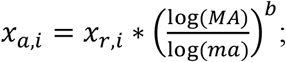

where *x*_*a,i*_ is the size adjusted lipid concentration for individual *i, x*_*r,i*_ is the raw lipid concentration for individual *i, MA* is the average mass for all individuals measured, *ma*_*i*_ is the mass for individual *i* and *b* is the slope of the relationship between the log of lipid concentration on the log of weight. The slope of the relationship was determined using linear regression of each log transformed lipid concentration against the log transformed fish weight across all sampled individuals. Size-adjusted traits (log transformed) were compared among reproductive stages and populations using ANOVA (Quinn and Keough 2002).

## Results

There was a relationship between log (lipid concentration) and log (body mass) (linear regression: F_1, 417_ = 83.65, p <<0.001) with a slope equal to 0.074658. I used this slope value to size-adjust individual lipid concentration.

There was an interaction between the reproductive stage and population location impacts on size-adjusted lipid concentration (F_8, 382_ = 2.859, p = 0.00425; Figure 2). Subsequently, I performed one-way ANOVA comparing lipid concentration among the reproductive stages for each population separately.

**Figure 2.**
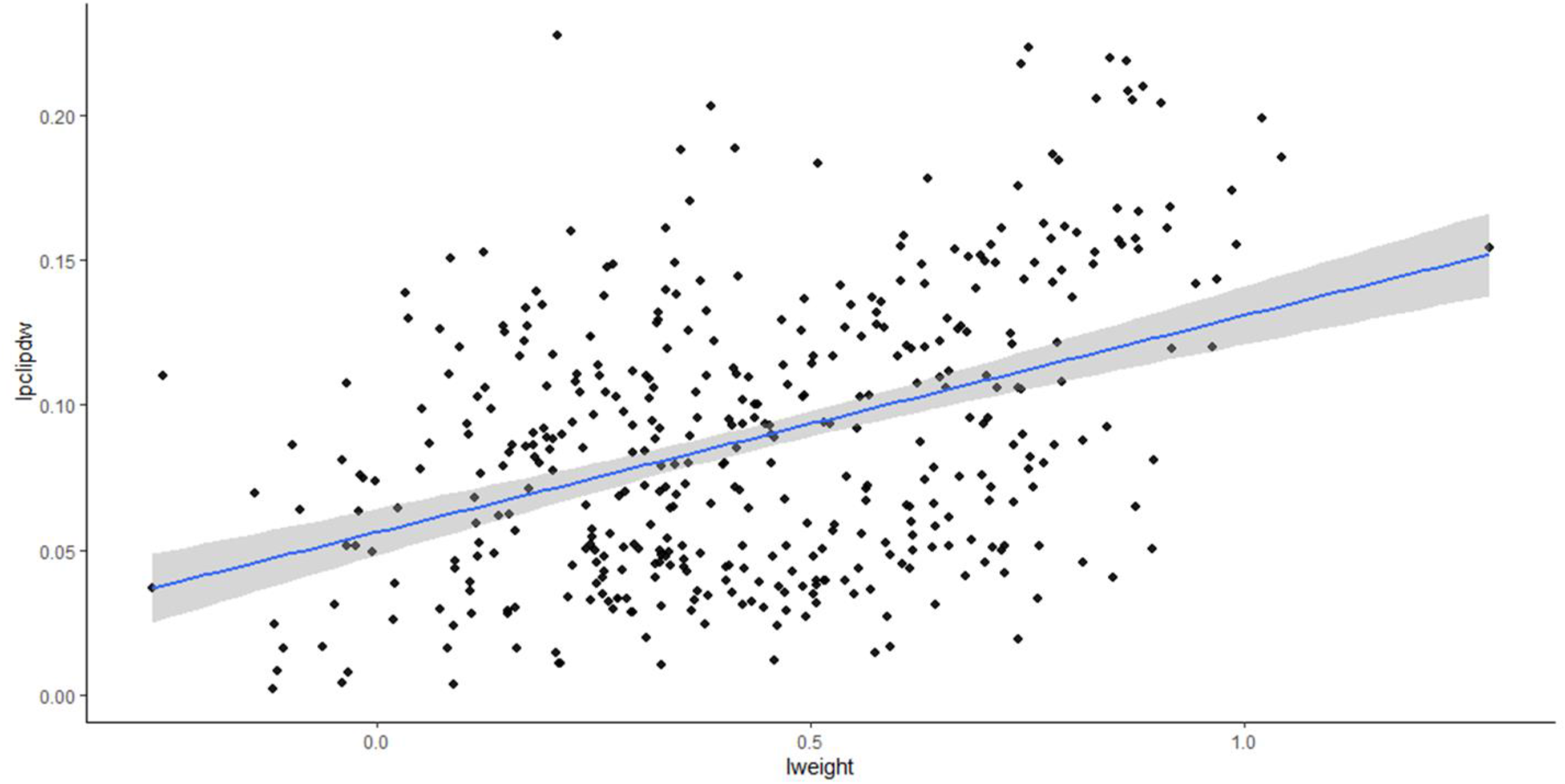
Scatter plot of log (lipid concentration) and log (weight) for all individuals sampled across the 5 locations with OLS regression line (with 95% CI – shaded region).

**Figure 3.**
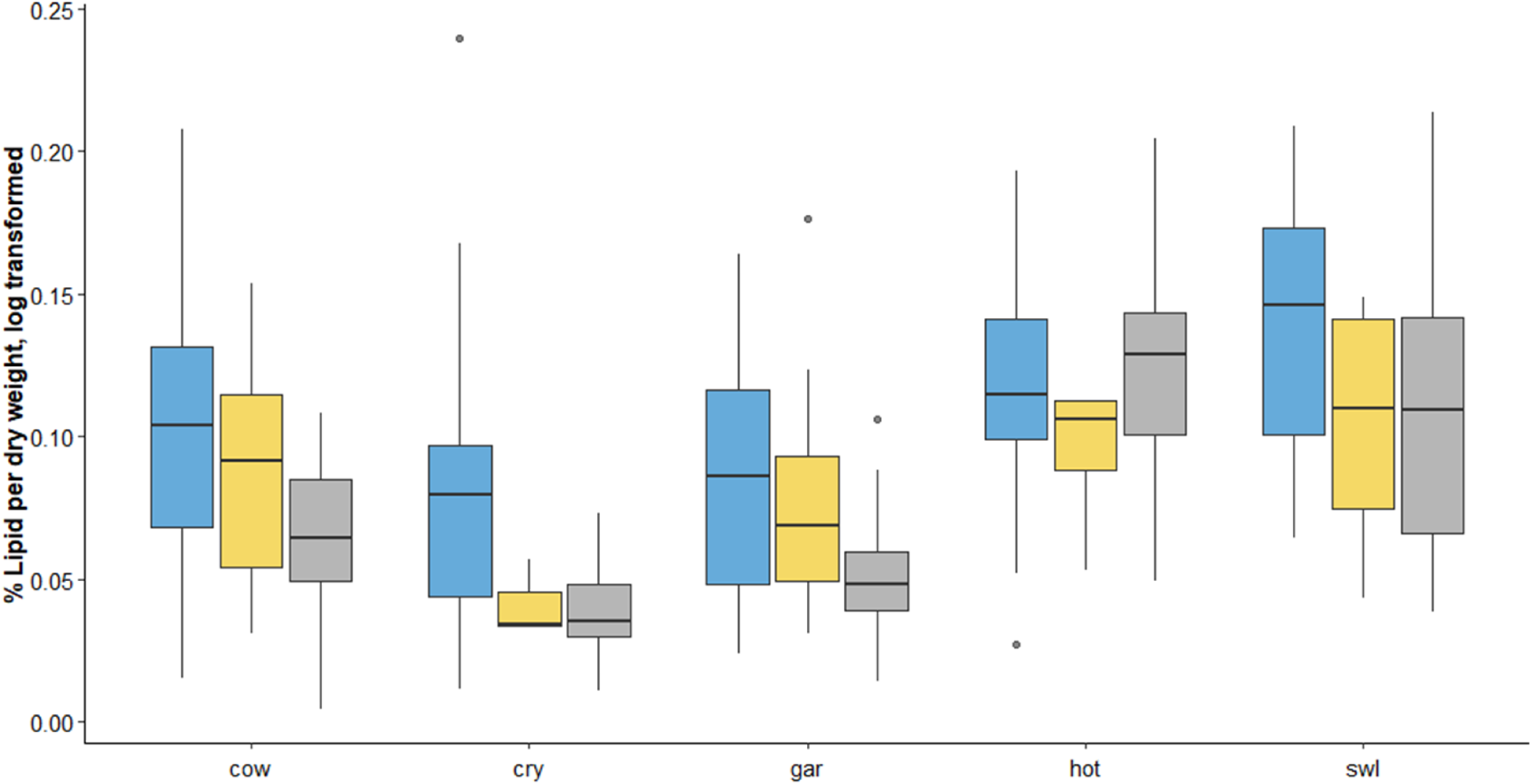
Boxplot of size adjusted lipid concentration (median – horizontal line; box – 1^st^ to 3^rd^ quartile range) for males sampled at courting (blue), parental care of embryos in the nest (yellow), and parental care of free-swimming young (gray) across the five locations sampled (cow – Cowichan Lake; cry – Crystal Lake; swy – Swy-a-Lana Lagoon; gar – Garden Bay Lake; hot – Hotel Lake). ANOVA results for each within population comparison among nesting intervals are indicated above each set of bars.

Lipid concentration decreased with progression of the reproductive interval for all populations except the population from Hotel Lake (Figure 2). For Cowichan Lake and Garden Bay Lake, lipid concentration decreased between the courting phase and early parental phase and decreased further from early parental to late parental phase. For Crystal Lake and Swy-a-Lana Lagoon, lipid concentration decreased from the courting phase to the early parental care phase but did not change between the early and late parental care phases.

### Discussion

Parental care activities are energetically expensive and intensity and duration of care is dependent on stored energy reserves when foraging opportunities during parental care are limited. In this study, I compared energy reserves (total lipid) of parental male threespine stickleback from two types of freshwater populations (limnetic specialists and benthic specialists) and a marine population (generalist ancestor to both specialist types; Bell and Foster 1993; Walker and Bell 2000; Hunt et al. 2008). As expected, lipid stores of parental stickleback from limnetic populations declined among the three intervals sampled; lipid concentration was highest during the courting interval, decreased between courting and care of embryos in the nest and further declined by the time males were caring for free-swimming embryos. Lipid stores also declined between courting and care of embryos in the nest for the males sampled from the marine population, but did not decrease further by the time males were caring for free-swimming embryos. Males from limnetic or marine populations tend to disengage from parental care once offspring are free-swimming. Males in limnetic populations will gradually begin another reproductive bout whereas males in marine populations will gradually move away from the nest site (Foster et al. 2008).

Lipid concentrations dropped in the limnetic and marine populations consistent with the assumption that foraging opportunities are limited during parental care. Cessation of parental care during the free-swimming embryo stage should lead to increased foraging opportunities and levelling off of decrease in lipid stores, as observed for the marine population. However, since the limnetic males likely began another reproductive bout their foraging opportunities remained low and their energy reserves continued to decline. Of the two benthic populations, lipid concentration of males in Hotel Lake did not change across the reproductive intervals as expected. However, in Crystal Lake, lipid concentration decreased more than for males in any other population. Though this is counter to what is expected for males in benthic populations, males in this population experience high level of conspecific nest predators (Foster cannibalism paper cites) and so it appears that the males in this population are expending a lot more energy on nest-defence than the are capable of recovering during opportunistic foraging opportunities during parental care, even though the foraging opportunities for males in this population may be high relative to the opportunities for nesting males in limnetic populations.

The stored energy (lipids) may have an impact on reproductive success for males in limnetic populations, and benthic populations that have high predation pressure. Energy reserves are necessary when foraging opportunities are limited. If energy reserves are inadequate, males terminate parental care before offspring are independent, or they may resort to filial cannibalism (FitzGerald 1991, 1992; Manica 2002). Female reproductive success in stickleback is tightly tied to the reproductive success of the males with which females leave their eggs. The challenge for females is to pick the best father, and in the case of limnetic sticklebacks, this would mean males with the highest lipid stores. Lipids mediate the uptake and transport of carotenoid pigments (Castenmiller and West 1998; Odeberg et al. 2003; Parker 1996; van het Hof et al. 2000; Yonekura and Nagao 2007) which make-up the striking red nuptial colouration displayed by male stickleback during reproduction (Olsen and Owens 1998; Wedekind et al. 1998). Hoelzer’s (1989) good parent hypothesis predicts that male signals should indicate male parental quality. Although Black et al. (2014) found a correlation between a carotenoid pigment (astaxanthin) and whole-body lipid, Scott and Black (2019) did not find a correlation between signal characteristics and stored lipid in the Hotel Lake population. Lack of correlation in Hotel Lake is not surprising since males in this population do not deplete lipids over the parental care interval so there is no need to signal lipids to prospective mates. On the other hand, I would expect a correlation between signal traits and lipid stores for stickleback in Garden Bay Lake. Correlation between stored lipid and male signal in Garden Bay Lake has not been examined.

In summary, I found that parental male stickleback from limnetic populations depleted lipids as the reproductive interval progresses. Stored lipids in these types of stickleback populations are clearly very important for fueling behaviours during parental care, and because stored lipid can influence female fitness should lead to a sexually selected signalling mechanism in these populations.

## Acknowledgements

Andre Laurin, Valerie Mucciarrelli, and Andonis Tzarougian assisted with collection of stickleback. Funding for the research was provided by an NSERC Discovery Grant. This research was reviewed and approved by the Animal Use Sub-committee of the Animal Care Committee at Western University, London, Ontario, Canada.

## Data availability

Data have been uploaded to Borealis and can be accessed at https://doi.org/10.5683 following publication of this manuscript in a per-reviewed journal.

## Ethics declaration

All field collections were carried out under

Western University Animal Care approval and under collection permits issued by the Province of British Columbia.

